# Supervised Capacity Preserving Mapping: A Clustering Guided Visualization Method for scRNAseq data

**DOI:** 10.1101/2021.06.18.448900

**Authors:** Zhiqian Zhai, Yu L. Lei, Rongrong Wang, Yuying Xie

## Abstract

The rapid development of scRNA-seq technologies enables us to explore the transcriptome at the cell level in a large scale. Recently, various computational methods have been developed to analyze the scR-NAseq data such as clustering and visualization. However, current visualization methods including t-SNE and UMAP are challenged by the limited accuracy of rendering the geometic relationship of populations with distinct functional states. Most visualization methods are unsupervised, leaving out information from the clustering results or given labels. This leads to the inaccurate depiction of the distances between the bona fide functional states and the variance of clusters. We present supCPM, a robust supervised visualization method, which separates different clusters, preserves global structure, and tracks the cluster variance. Compared with six visualization methods using synthetic and real data sets, supCPM shows improved performance than other methods in preserving the global geometric structure and data variance. Overall, supCPM provides an enhanced visualization pipeline to assist the interpretation of functional transition and accurately depict population segregation.

## 1 Introduction

Single-cell RNA sequencing (scRNAseq) technology represents rapidly expanding opportunities to interrogate cellular functions at an unprecedented resolution (Sandberg, 2014; Macosko *et al*., 2015; Ziegenhain *et al*., 2017). In the meantime, numerous computational methods have been developed for scRNA-seq data sets among which clustering and visualization are the two most important and related procedures (Chen *et al*., 2019; Kiselev *et al*., 2019). Various kinds of clustering algorithms are introduced, such as PhenoGraph (Levine *et al*., 2015), Seurat (Hao *et al*., 2020), SC3 (Kiselev *et al*., 2017), scanpy (Wolf *et al*., 2018), CMDS (Little *et al*., 2018). As for visualization, a main challenge is due to the fact that the higher dimensional body has a substantially larger capacity than what is visually effective on a two-dimensional space. This frequently results in overlapping and crowding of of the embedded data points, which is also known as crowding issue. To overcome this challenge, two nonlinear methods, t-SNE and UMAP, were developed and have been widely adopted in the current scRNA-seq toolkits (Linderman *et al*., 2019; Zhou and Jin, 2020; Becht *et al*., 2019). These methods fit Gaussian distributions with heavy-tailed distributions, creating “repulsive forces” to avoid the overlapping. However, due to the difficulty in controlling the strength of the force, these methods tend to overly stretch the data with unnecessary information loss. Specifically, t-SNE minimizes KL-divergence between Gaussian and t-distributions derived from the pairwise distances in the high- and low-dimensions, respectively (van der Maaten and Hinton, 2008). Although t-SNE is effective in preserving the local structure, it cannot precisely render the global structure. For example, two points that are far apart in the original high-dimensional space may be close under the t-SNE coordinates or vice versa. With cross-entropy instead of KL-divergence, UMAP can preserve more large-scale structure compared to the t-SNE (Becht *et al*., 2019). Despite their popularity, UMAP and t-SNE have several known limitations (Wattenberg *et al*., 2016). As mentioned above, often times, the inter-cluster distances, or the long range distance, are not meaningful as the consequence of their adoption of the conditional probability. Two clusters close under UMAP or t-SNE coordinates maybe be far apart in the original dimensions, which introduces challenges in populations with transitional functional states. Furthermore, the variance of clusters in UMAP and t-SNE does not correlate with the actual variance. Instead, it is largely driven by the sample size.

A main thrust of the current scRNA-seq analysis pipeline is the clustering algorithms that can produce high-quality class labels, which are an untapped source of information to improve the current visualization methods. Therefore, we should use this piece of information to assist the visualization if possible. Because although visualization methods such t-SNE and UMAP also enforce formation of clusters, their ability to detect labels is suboptimal, since as visualization methods, they need to map the data to extremely low dimensions in which the clusters are less separated (Kiselev *et al*., 2017). For example, if we perform a k-means clustering on the two dimensional UMAP embedding of the RNAmix data with 7 clusters (Fig. 1), a large portion of cell labels will be incorrect. In contrast, 94.1% of the cluster labels generated by Seurat match with the underlying cell types. This raises an important question: how to incorporate the almost perfect class labeling information returned by a pure clustering algorithm into the visualization result? In other words, what is the best way to design supervised visualization techniques?

**Figure 1:**
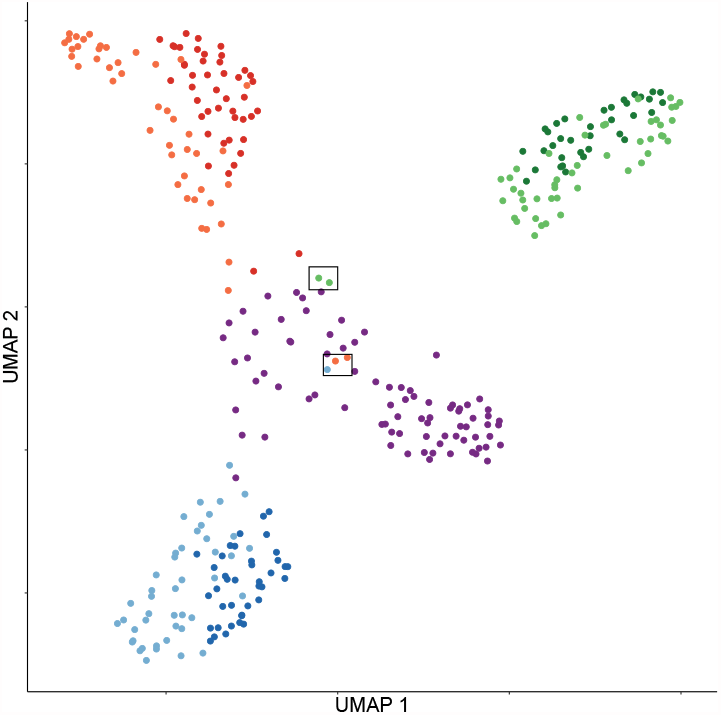
UMAP visualization for RNAmix data. Cells are colored according to labels from clustering algorithm in Seurat. Highlighted cells suggest the conflicting result between clustering and visualization methods.

There are only a few supervised visualization algorithms available in the literature, such as supPCA (Barshan *et al*., 2011) and supUMAP (McInnes and Healy, 2018). However, supPCA is a linear method that is unsuitable for scRNAseq data where the transcriptome data is nonlinear. Compared to the vanilla UMAP, supUMAP utilizes label information to debond “wrong” neighbors that actually belong to different clusters. However, since the vanilla UMAP emphasizes much more on the formation of clusters than the preservation of geometry, it has similar advantages and disadvantages to clustering methods and hence adding label information to it has relatively small improvements on its result (see results in section 3). We argue that it would be more beneficial to supervise the geometry-preserving visualization methods such as PCA, LLE (Roweis and Saul, 2000), MDS, etc, because those methods have complementary merits to clustering algorithms and combining them may give rise to an algorithm having the merits of both.

One major issue with these classical geometry-preserving methods is the aforementioned crowding issue. Recently, Wang and Zhang (2019) proposed a novel visualization algorithm, Capacity Preserved Mapping (CPM) to address it, therefore can better preserve the geometry. However, CPM alone does not enforce well-separated clusters to appear in the visualization, which sometimes cause it hard to digest. This drawback may be compensated using the known label information. In this paper, we developed a supervised CPM (supCPM) algorithm which runs the vanilla CPM for the first phase and in the second phase optimizes the objective function with an extra supervised term. We demonstrate the superior performance of supCPM over other visualization algorithms on both synthetic and real data sets.

## 2 Methods

### 2.1 Relative Capacity

Suppose **X**_*n*× *p*_ = (*x*_1_, …, *x*_*n*_)^*T*^ ∈ ℝ^*n×p*^ is the RNA-seq data set with *n* cells and *p* genes. From manifold learning’s perspective, people commonly assume the high-dimensional data, *x*_*i*_, live on a space with a small intrinsic dimension since genes are closely related with each other. The small intrinsic dimension assumption ensures that the data can be projected to the low dimensions for visualization. More rigorously, we assume that each data point is independently drawn at random according to a sampling probability *f* from some underlying low-dimensional compact manifold ℳ living in the high dimensional space ℝ^*p*^. Suppose *x* ∈ ℳ, the relative capacity of a radius-*r* neighborhood centered at *x* is defined as

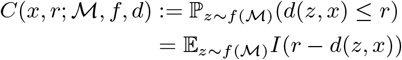

where *d* is the pairwise distance defined on ℳ and *I*(·) is the indicator function. The relative capacity could be intuitively interpreted as the expected number of points in a neighborhood.

Further, the average capacity of the radius *r* neighbourhoods is

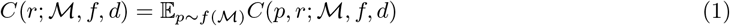

that is the averaged relative capacity across all the locations. By the definition, it’s easy to check that functions *C*(*r*; ℳ, *f, d*) is a cumulative distribution, and we define

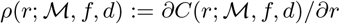

which is a density function.

### 2.2 Capacity Adjusted Distance

Capacity preserved mapping aims at preserving the relative capacity defined in (1) at all scales *r*. The preservation of capacity is crucial to avoid crowding issue mentioned above.

To avoid the crowding issue at a minimal cost, Wang and Zhang (2019) defined the Capacity Adjusted Distance (CAD)

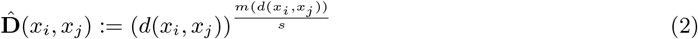

where *d*(·, ·) is a user chosen distance (Eulidean, geodesic, diffusion, etc), *s* ∈ {2, 3} is the visualization dimension, *m*(*r*) is the intrinsic dimension of the data at scale *r* that can be estimated from the data (Wang and Zhang, 2019).

CAD is shown to be preservable under dimension reduction without introducing crowding. Namely, there exists a map *P*, such that *C*(*r*; ℳ, *f, d*) = *C*(*r*; 𝒮, *f*_uniform_, ‖ ·‖_2_), where 𝒮 = *P* (ℳ) is the embedded data manifold equipped with the Euclidean metric ‖ ·‖_2_ and the uniform sampling distribution. That is to say, the data density did not change during embedding.

In practice, we slightly modify 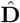 to ensure that the ranking of the pairwise distances under 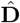 is consistent with that under the user input distance *d*(·, ·). Let 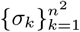 be the set of all the pairwise distances of original data under *d*(·, ·) in ascending order with 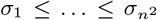. The modified Capacity Adjusted Distance 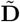 is then defined recursively for *k* = 1, …, *n*^2^ as 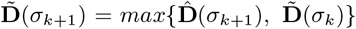. Then 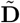 is monotonously nondecreasing 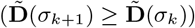, meaning that larger distances are still larger after the capacity adjustment. Therefore, the relative distance is preserved.

### 2.3 Capacity Preserved Mapping

Let 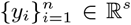 be the low dimensional embedding coordinates for each cell and **Y** = (*y*_*i*_, …, *y*_*n*_)^*T*^ be the matrix representation. From scRNAseq data set **X**_*n×p*_, we compute the modified capacity adjusted distance matrix 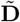. The pairwise distances in both high and embedding spaces are changed to probability distributions by taking a reciprocal (a small positive constant number is added to avoid taking the reciprocal of 0) followed by a normalization. One minimizes Kullback-Leibler divergence to get the embedding points 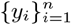. That is to say

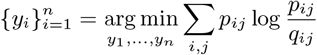

where,

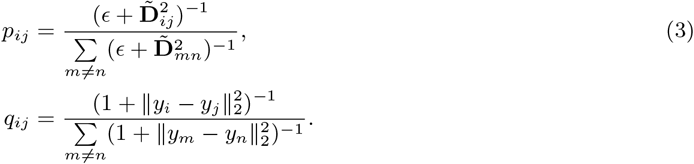

A major difference from t-SNE is that the above probabilities *q*_*ij*_ and *p*_*ij*_ in CPM are not conditional probabilities, therefore reducing information loss.

### 2.4 Supervised CPM

The ability of CPM of displaying global geometric information makes it a good complement of various clustering algorithms. This is the motivation of Supervised CPM, where we hope to integrate the label information into CPM without too much distortion of the preserved geometry.

Supervised CPM has two modifications upon CPM. First, a second term containing the label information is added to the objective

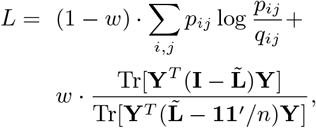

where *w* is a scalar turning parameter and 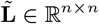 is the label matrix defined as 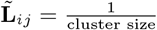 if *i*-th and *j*-th points have the same label and 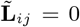 otherwise. The trace ratio is a measure of the cluster separation (Liu *et al*., 2013; Jia *et al*., 2009). Specifically, the numerator measures the inter-cluster distance and the denominator is the intra-cluster distance, therefore maximizing the ratio means maximizing the cluster separation. When solving this non-convex optimization, we use the result of the vanilla CPM as an initial guess to avoid being trapped at local minimizers.

The second change to CPM is that we modify the distances 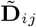 in such as way that the inter-cluster distances are magnified by a constant *C*(*C* ≥ 1) before inserting into (3) to define the probability. This modification is particularly useful when the data is contaminated by large noise. Usually, the metric used by clustering algorithms ^*^ is more robust to noise than the Euclidean or geodesic distances used in CPM. When the large noise cause clusters to overlap under the metric of CPM, magnifying the inter-cluster distances would of great help. We comment that this step cannot replace the usage of the trace ratio objective because 1) scaling all inter-cluster distances by one scaling factor is very crude 2) the magnification factor *C >* 1 should be small (say, smaller than 2), otherwise it may increase the dynamic range of pairwise distances by too much for CPM to preserve ^†^. We choose *C* = 1.3 in practice. For clean data, this modification is not necessary.

To avoid the deformation of clusters’ shape during supervised optimization, we also set the degree of freedom of high dimensional t-distribution from 1 to *d*, a customized parameter. That is to say,

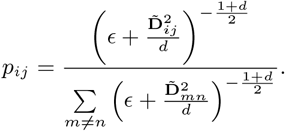

A larger degree of freedom means longer distance will be enlarged further. Thus, increasing the parameter *d* will weaken inter-cluster influence so that during the supervised phase, the shape of cluster will not shrink asymmetrically. But *d* shouldn’t be too large as well, since the mismatch of probability distribution in both high and low dimensions could cause information lost. More details for the effect of different values of *d* on the embedding results can be found in the Supplementary Material.

As for the choice of distance, we recommend to use Euclidean distance for noisy data set, and geodesic distance for clean data set. The detailed steps are summarized in Algorithm 1.

#### Algorithm 1: supCPM

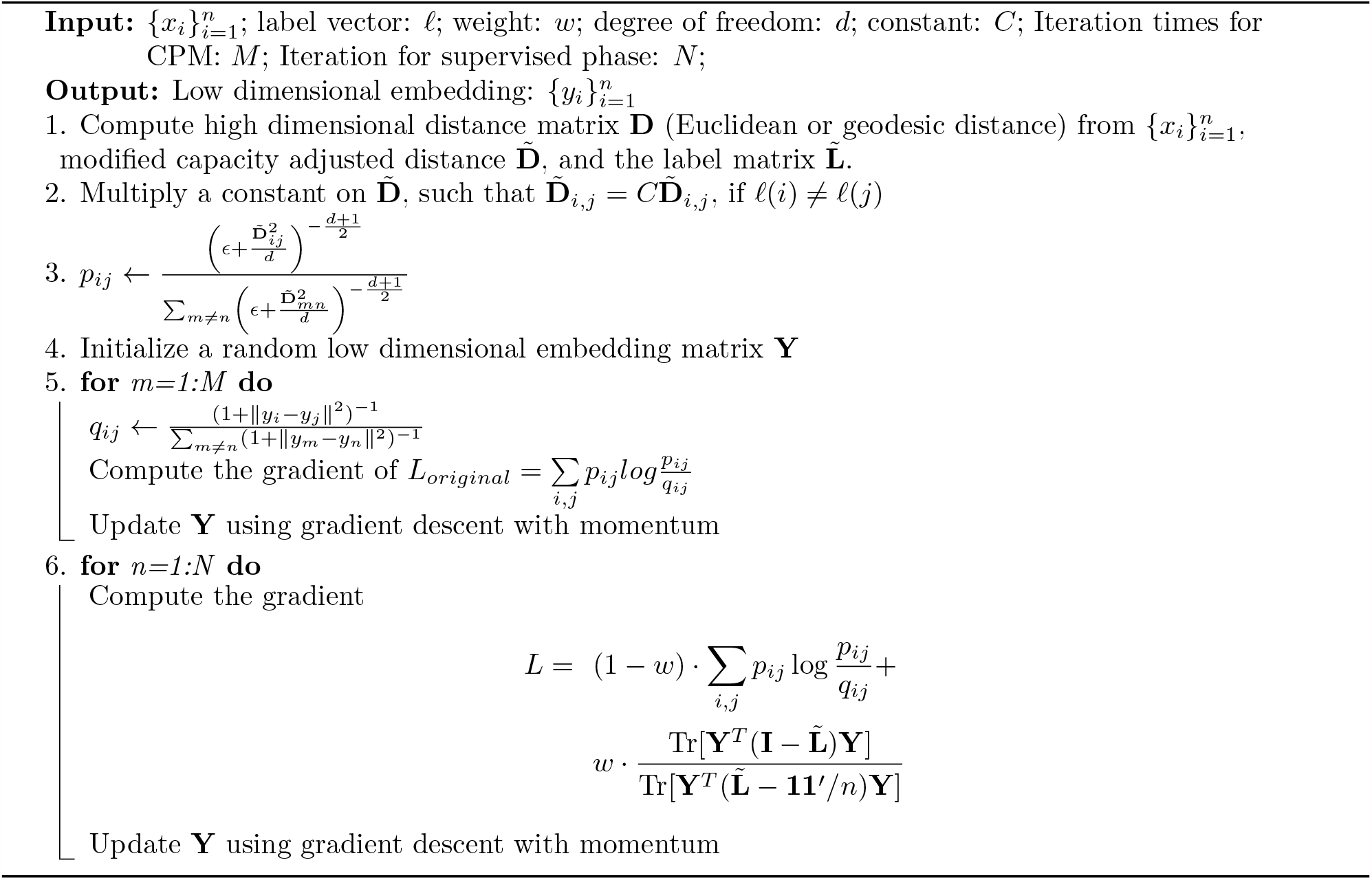

#### 2.5 Evaluation Metrics

Following Kobak and Berens (2019), we use five metrics to evaluate the preservation of both geometry structure and cluster variance, and separation of clusters for quality comparison purpose.

##### KNC

The fraction of <u>**k**</u>-<u>**N**</u>earest neighbors of <u>**C**</u>luster means in the high dimensional data which are still preserved as the *k*-nearest cluster means in the embedding space, that is to say,

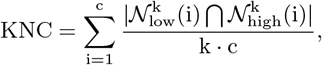

where 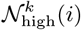 and 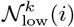 are the sets of k-nearest neighbors of the *i*-th cluster mean in high and low dimensions respectively, and *c* is the number of clusters. KNC depicts the proportion of cluster centers preserved. Intuitively, it captures the mesoscopic structure and measures whether the relative positions of clusters are projected correctly.

##### KNN

The fraction of <u>*k*</u>-<u>**N**</u>earest <u>**N**</u>eighbor pairs in the high dimensional data that are still *k*-nearest neighbor pairs in the embedding dimensions (Lee and Verleysen, 2009). Specifically, we define

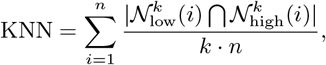

where 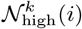 and 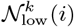 are the sets of *k*-nearest neighbors of *i*-th cell, and *n* is the sample size. KNN captures the local connectivity, or microscopic structure, preserved by the visualization methods.

##### CPD

Spearman <u>**C**</u>orrelation between <u>**P**</u>airwise <u>**D**</u>istances in the high and low dimensions, is defined as follows (Becht *et al*., 2019):

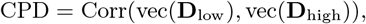

where **D**_low_ and **D**_high_ are the pairwise distance matrices in the low and the high dimensional space. vec(·) creates a column vector from a matrix by stacking its columns. CPD provides an evaluation of the visualization methods in a global, macroscopic view.

##### CorV

Spearman <u>**Cor**</u>relation between within-cluster total <u>**V**</u>ariation in the high-dimensional space and those in the low. It measures the relative variation of each clusters preserved by dimensional reduction methods. Suppose that *X* and *Y* are data matrices of high and low dimensions respectively, *V*_**X**_ and *V*_**Y**_ are vectors of variations of each cluster with *V*_**X**_(*i*) = Tr(Cov(**X**_i_)), **X**_*i*_ is the coordinate matrix of the *i*-th cluster. *V*_**Y**_ is defined similarly. CorV is then calculated by

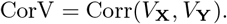

Larger values of CorV means a better preservation of cluster radii.

##### LogFisher

Measurement of how well the visualization algorithms separate clusters. It is defined as

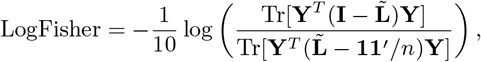

**Y** is the embedded data matrix and 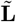 is the label matrix previously defined. A higher logFisher score indicates clusters are more separated.

### 3 Results and Conclusions

#### 3.1 Synthetic Data set

Following the procedure in Kobak and Berens (2019), we simulated 3600 points from 9 different multivariate Gaussian distributions in 50 dimensions. The data can be grouped into three main classes, and each class can further be separated into three sub-clusters. Data were sampled from Gaussian distributions with different covariance matrices, which were **I**, 1.5^2^**I**, and 2^2^**I** for sub-clusters in the first, the second and the third classes respectively, where **I** was the identity matrix. Points from different classes were then shifted by 20 in mutually orthogonal directions. Next, all points from each sub-cluster in the first class (n = 600 per type) were additionally shifted by 5 in mutually orthogonal directions; by 10 in the second class (n = 400 per type); and by 15 in the third (n = 200 per type). In addition, we made the directions differentiating sub-clusters orthogonal to the directions differentiating the three main classes. To explore the sample size effect among different methods, we specifically let sub-clusters with larger sample size had smaller variation. For comparison, we visualized the simulated data using Multidimentional Scaling (MDS), PCA, supPCA, t-SNE, UMAP, supUMAP, CPM, and supCPM as shown in Fig 2.

**Figure 2:**
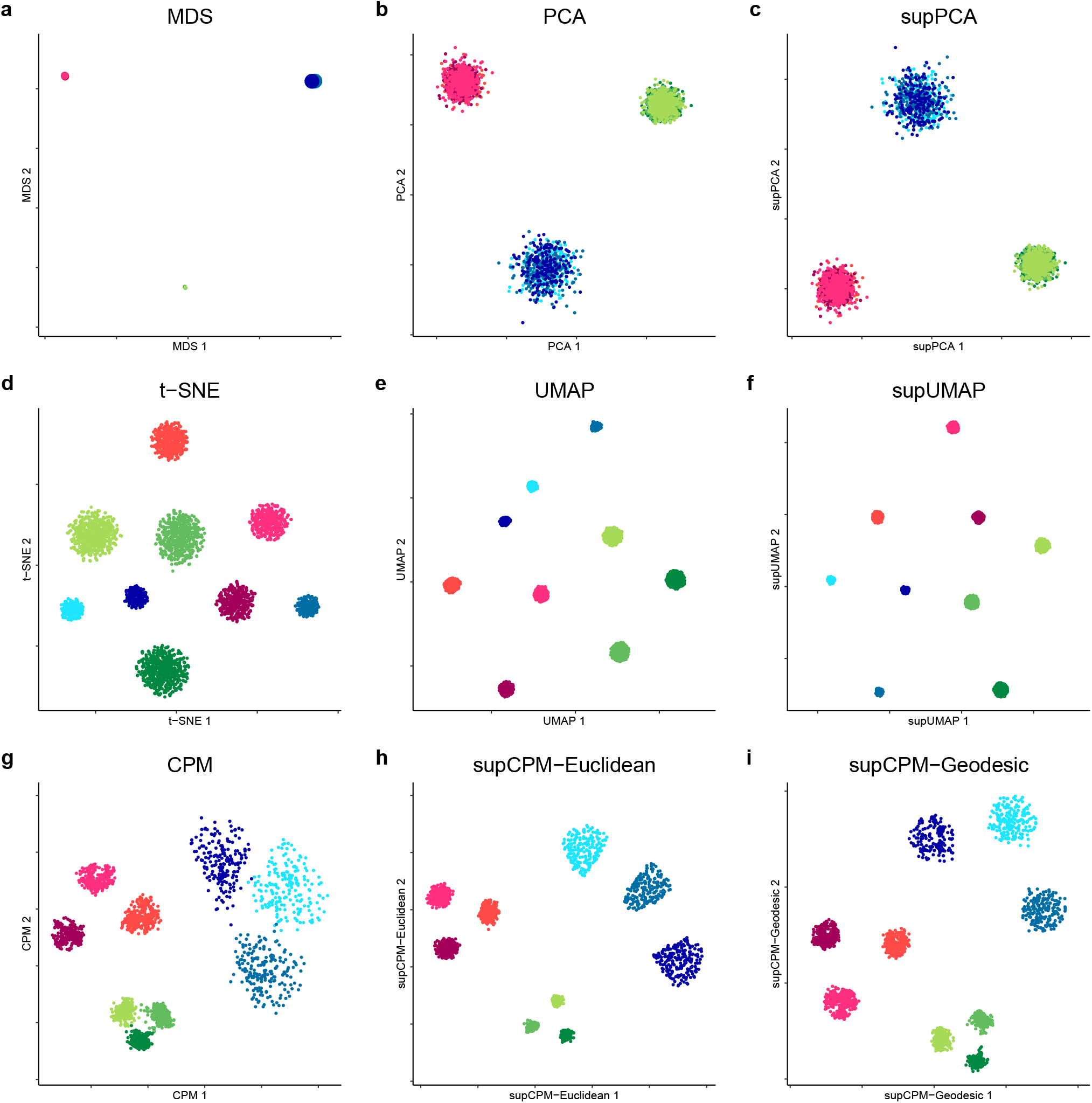
Visualization of Synthetic data. (**a**) MDS on cluster means. The size of point is proportion to the variance of cluster. (**b**) PCA. (**c**) supervised PCA.(**d**) t-SNE. (**e**) UMAP with cosine distance. (**f**) supUMAP with cosine distance. (**g**) CPM. (**h**) supCPM with Euclidean distance. (**i**) supCPM with Geodesic distance.

In Fig 2a, MDS was applied on cluster means instead of individual data, and the size of the resulting six points was proportional to the true variance per cluster in the original data. The MDS results matched well with the global geometric structure of sub-clusters. In Fig 2b-2c, PCA and supPCA captured the variance of clusters well, but clusters were indistinguishable because the sub-cluster directions are orthogonal to the first two PC directions. As shown in Fig 2d-2f, t-SNE, UMAP and supUMAP distorted the variance of clusters and showed that the three green sub-clusters have the largest variance. This is because the green sub-clusters had the largest number of points. The metric, CorV, in Fig 6 reflected the same observation. Apart from the issue of preserving variance, t-SNE and UMAP severely distorted the data geometry, with t-SNE exhibting a worse performance as shown in Fig 2d and 2e. In contrast, CPM and supCPM showed desirable visualization results (Fig 2g - 2i). Even though a small fraction of points in green clusters were mixed up, CPM successfully maintained the variance and geometry structure. Similarly, supCPM not only separated different clusters more effectively, but also kept the most of geometry and variance structures. Notably, supCPM with geodesic distance also kept the isometric structure with high fidelity. In addition, the KNN metric also demonstrated that most of the local structure was also precisely rendered by supCPM. Finally, based on Fig 6, supCPM achieved high scores in all evaluation metrics, outperforming the current benchmarks.

#### 3.2 scRNAseq Data sets

##### 3.2.1 Synthetic scRNAseq Data

We used the synthetic scRNAseq, denoted as RNAmix, generated by Tian *et al*. (2019) that contained 340 ‘pseudo cells’. These ‘pseudo cells’ with various ratios of RNA contents extracted from three cell lines (H2228, H1975, HCC827) formed seven clusters with a triangle structure (Fig. 3a). Owing to the small sample size, we performed log-Normalization, scale and PCA on the data following the standard Seurat procedure (Hao *et al*., 2020). We then took the top 10 PCs as the input for both the shared nearest neighbor (SNN) modularity optimization-based clustering and the visualization methods in Seurat. The SNN method successfully grouped the cells into seven groups reflecting the true underlying structure with an error rate of 5.9%. Because in practice we rarely know the true cluster labels, the resulting SNN estimated cluster labels were used as input for the three supervised visualization methods.

**Figure 3:**
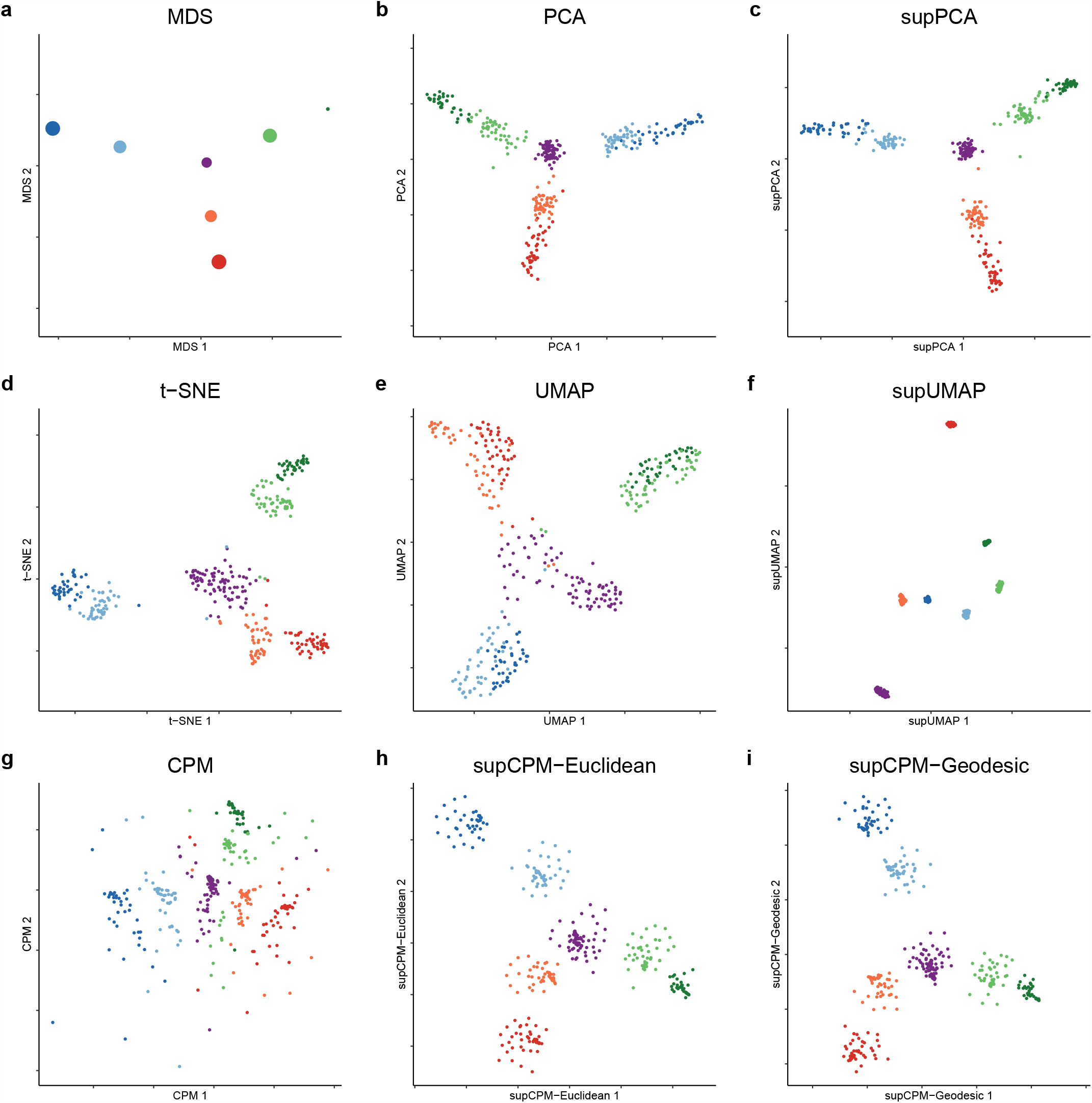
Visualization of RNAmix data. (**a**) MDS on the cluster means. The size of point is proportional to the variance of cluster (**b**) PCA. (**c**)supPCA. (**d**) t-SNE. (**e**) UMAP with cosine distance. (**f**) supUMAP with cosine distance. (**g**) CPM. (**h**) supCPM with Euclidean distance.(**i**) supCPM with geodesic distance.

As the global structure of this data set is linear, PCA and supPCA performed well in uncovering the triangular structure but failed to effectively separate the clusters as shown in Fig 3b-3c. Fig 3d shows that t-SNE preserved some global structure, but the triangle structure was blurred. UMAP distorted the structure even further. In particular, in Fig 3e, it was challenging to ascertain the underlying affinity between the central purple cluster with the surrounding units. CPM produced a few floating points due to the high noise level of the data set but captured the correct triangle structure. In contrast, by utilizing the cluster labels, supCPM forced those floating cells to be closer with other cells in the same cluster and hence generated better results reflecting the true variance and global structure than the other methods. Furthermore, supCPM respected the local structure at the same level as UMAP as summarized in Fig 6.

##### 3.2.2 Cell Line Data

To evaluate the performance of supCPM in real data sets, we downloaded and analyzed the expression profiles of cancer cells lines generated by McFarland *et al*. (2020). The whole data set contains cells from twenty-four cell lines treated with either Nutlin or vehicle control dimethyl sulfoxide (DMSO). For the ease of visualization, we chose eleven cell lines with 1224 cells under the DMSO treatment. The selected cell lines were originated from six of organs: RCC10RGB (Kidney), LNCAPCLONEFGC (Prostate), BICR31 (Upper Aerodigestive Tract 1), BICR6 (Upper Aerodigestive Tract 2), SQ1 (Lung 1), RERFLCAD1 (Lung 2), NCIH2347 (Lung 3), CCFSTTG1 (Nervous System 1), DKMG (Nervous System 2), RCM1 (Large Intestine 1), LS1034 (Large Intestine 2). We used Seurat SCTransformation to normalize and scale the data before applying PCA. The top 20 PCs were used for clustering to recover the underlying cell lines structure, which resulted an accuracy of 99.9%. We also annotate each cluster based on the known cell line labels.

Cell lines derived from the same organ tended to have similar expression profiles and hence should be plotted close to each other. This is indeed the case as in shown the MDS result (Fig 4a). Among the rest of the methods, only CPM and supCPM displayed the cell lines correctly reflecting their origins, such as the Lung, large intestine, upper aerodigestive tracts, and the nervous system. Also, we observed that the prostate cancer cells behaves differently from the other cell lines, which was also captured by supCPM and CPM. Regarding to numerical metrics as in Fig 6, supCPM outperformed t-SNE and (sup-) UMAP in KNC, CPD and CorV. supCPM showed a comparable performance as UMAP on the local metric KNN. In sum, supCPM preserves substantially more global structure and variance than t-SNE and UMAP while keeping the local structure similar as UMAP.

**Figure 4:**
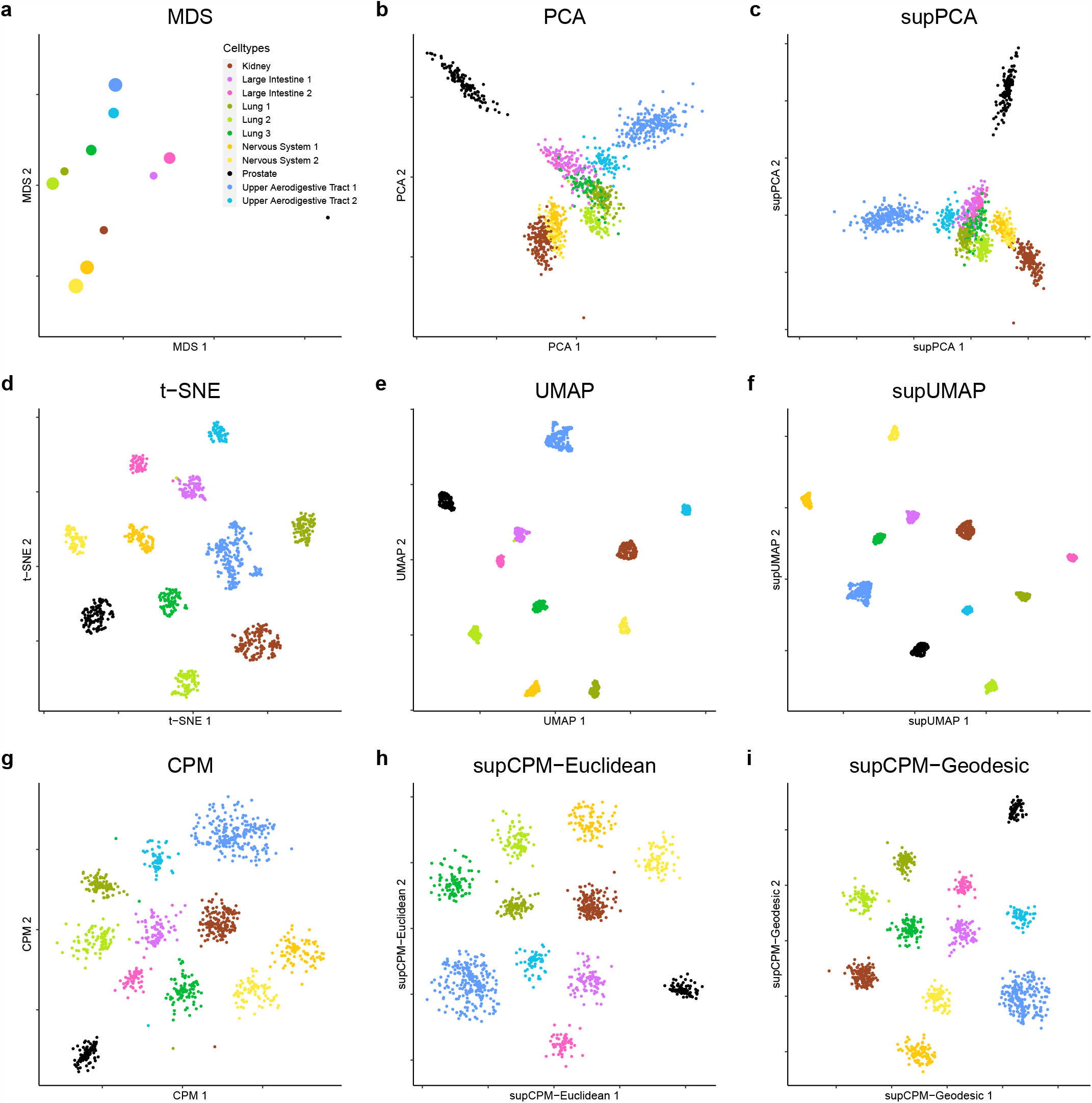
11 Cancer cell lines data. (**a**) MDS on cluster means. The size of point is proportional to the variance of cluster. (**b**) PCA. (**c**) supPCA. (**d**) t-SNE. (**e**) UMAP with cosine distance. (**f**) supUMAP with cosine distance. (**g**) CPM. (**h**) supCPM with Euclidean distance. (**i**) supCPM with geodesic distance.

##### 3.2.3 Human Peripheral Blood Mononuclear Cell Data

In addition to cell line data, we next applied supCPM to 2700 human peripheral blood mononuclear cells (PBMC) data derived from a healthy donor, which was downloaded from the 10x Genomics website. Following the standard Seurat procedure, we clustered the cells into twelves groups and annotated them using predefined marker genes. Platelets do not have nuclei and are not considered true cells. They are cellular fragments with a greatly different gene expression profile from the other hematopoietic cellular subsets. Thus, platelets ought to be far away from the other cells as the result in Fig 5a. However, neither t-SNE nor UMAP revealed this information (Fig 5d-5f). In contrast, SupCPM successfully recapitulated this structure. In addition, dendritic cells (DC), CD14^+^ Monocyte (Mono) and FCGR3A^+^Mono are myeloid cells and should differ from the lymphoid compartment. Results from supCPM clearly grouped the three myeloid subsets together, while the other methods have misleading results. In particular, UMAP placed B cells close to DC cells instead of other lymphoid lineage cells. As evidence of visualizing the true biologic relationships among clusters, we also focused on memory CD4^+^ T-cells and naive CD4^+^ T-cells. UMAP and t-SNE almost projected these cell types with the largest variation, while the truth showed in MDS was that these two types had very small variations. In contrast, supCPM successfully preserved the biologic relationship and projected the structure onto a 2D space. In sum, both visualization figures and evaluation metrics support that supCPM shows an unprecedented rigor in preserving the geometry and variance information of complex cellular architectures.

**Figure 5:**
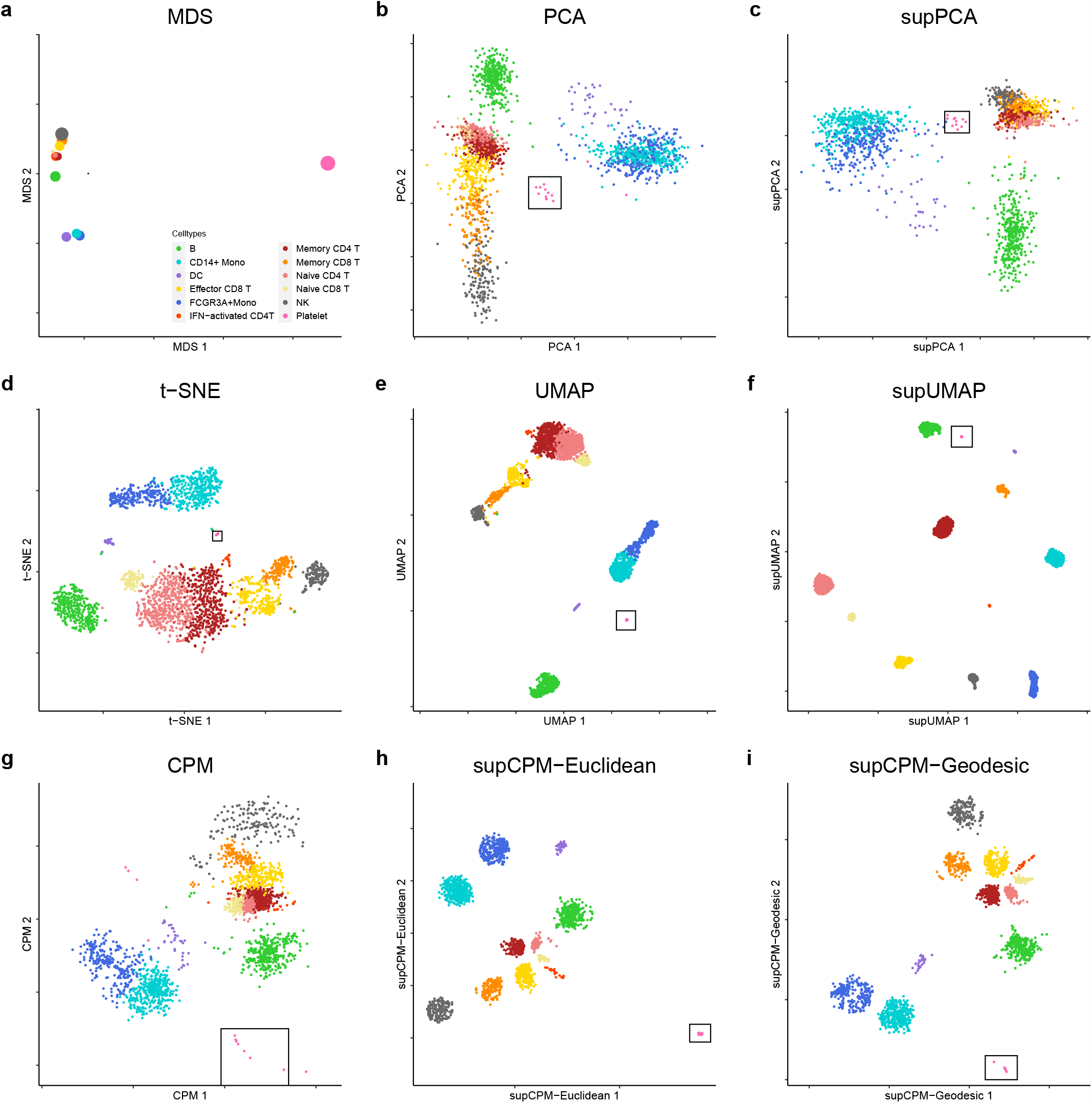
PBMC data. (**a**) MDS on cluster means. The size of point is proportional to the variance of cluster. (**b**) PCA. (**c**) supPCA. (**d**) t-SNE. (**e**) UMAP with cosine distance. (**f**) supUMAP with cosine distance. (**g**) CPM. (**h**) supCPM with Euclidean distance. (**i**) supCPM with geodesic distance. Platelet cells are highlighted in a black box in figures

**Figure 6:**
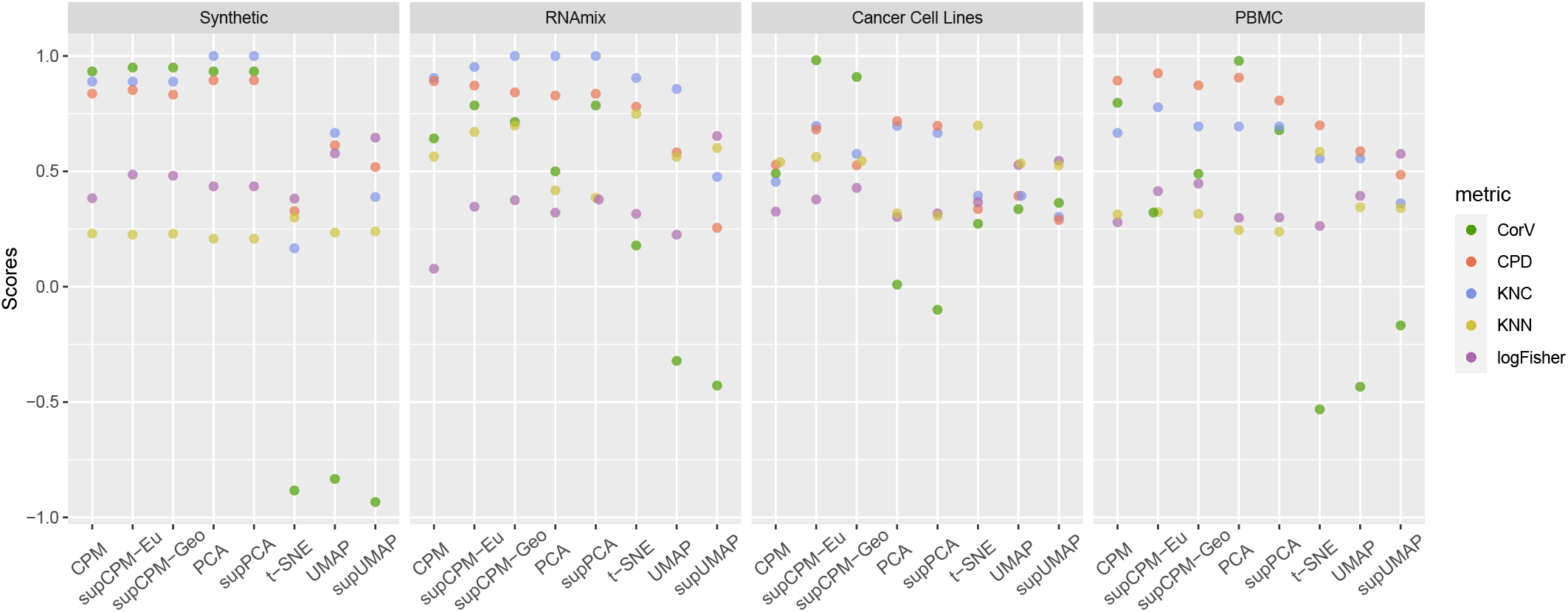
Visualization methods Comparison using five metrics. Some points are jittered hori-zontally to avoid overlap.

### 4 Discussions

The rapidly expanding need to reveal the functional relationships at single-cell resolution poses important challenges to the current visualization tools, which do not always render the bona fide cluster segregation in a 2D space. Thus, a more accurate and functionally related single-cell profiling pipeline is in urgent need. In this study, we propose supCPM, a supervised visualization method, to visualize the true functional relationships between cell types by leveraging both geometry information and label information. The implementation of supCPM is timely because of the frequent inconsistency between the visualization and clustering results. Not only does supCPM integrate the label information, but it also substantially improves the visualization fidelity and shows superior performances over the current benchmarks. And surprisingly, even though supCPM preserves geometry globally, it still preserves local structure at the same level as UMAP. Another major limiation of UMAP and t-SNE is that both methods cannot preserve the variation of clusters. The variation of clusters is largely associated with the sample size rather than the true variation. supCPM alleviates the problem and makes the visualization more functionally relevant. To test the performance of supCPM, we compared it with six other visualizing algorithms and evaluated them by five metrics. Metric results together with the visualization figures substantiates the unprecedented rigor of supCPM in recovering the true variation, keeping both global and local structures, and distinguishing different cell types.

Accurate interpretations of single-cell data sets depend on effective visualization. supCPM is an example of leveraging an exteneded array of information. There are additional possibilities to further improve the visualization algorithms. Firstly, how to combine the intermediate structure of clustering algorithm could be a meaningful question. supCPM only takes the final labels and leaves other information out. For instance, the SNN graph produced during Phenograph clustering could provide more information for visualization, so that the visualization could be more consistent with the underlying clustering algorithm. Secondly, there is still other relevant information could be utilized. Tree structure indicating the affinity of cell types could be helpful for visualization. Tree structure showing which two clusters are closer to each other will help the visualization to keep global structure. It would be helpful to incorporate these information to increase the accuracy and interpretability. We only tested supCPM on data sets with discrete cell types. Processing the label information from complex scRNAseq data sets with continuous functional transition may be challenging. Continuous scRNAseq data often exhibits trajectories where functional overlaps occur. This real world challenge could limit effectiveness of supCPM, because the second optimization part separates different clusters far apart. The solution to this problem entails a more delicate supervision method. And more difficultly, one could think of how to process the data set with the mixture of both discrete and continuous cell types. Manual review of the functionally overlapping clusters with known gene matrices that inform distinct lineage subsets would be equally important. Overall, here we present a novel and rigorous new visualization algorithm, which shows accurate representation of underlying functional relationship and outperforms the current benchmarks.

## Acknowledgements

We would like to acknowledge Leland McInnes for his feedback and explanations and Ali Ghodsi for the code of supervised PCA. This work was supported in part by the NIH grants U01DE029255 and RO3DE027399 and the NSF grants DMS-1902906 and NCS-1630982.

## Data Availability

Raw data of RNAmix are available at GEO under accession code GSE118767, and the processed data are available at https://github.com/LuyiTian/CellBenchdata. The Cancer cell line data are available at https://figshare.com/s/139f64b495dea9d88c70. The PBMC data set is available at 10x Genomics’s website https://support.10xgenomics.com/single-cell-gene-expression/datasets.

## Code Availability

Code used for supCPM, comparison analysis and figure generation is availalbe at https://github.com/zhiqianZ/supCPM.git.

## Author Contributions

**Conceptualization:** Zhiqian Zhai, Rongrong Wang, and Yuying Xie.

**Data curation:** Zhiqian Zhai.

**Formal analyses:** Zhiqian Zhai.

**Methodology:** Zhiqian Zhai, Rongrong Wang, and Yuying Xie.

**Project administration:** Rongrong Wang and Yuying Xie.

**Resources:** Yu L. Lei, Rongrong Wang, and Yuying Xie.

**Software:** Zhiqian Zhai.

**Writing-original draft:** Zhiqian Zhai.

**Writing-review & editing:** Yu L. Lei, Rongrong Wang, and Yuying Xie.

## Competing interests

The authors declare no competing interests.

## Footnotes

* For example, cell similarity is measured with the proportion of shared nearest neighbors in the clustering methods PhenoGraph and Seurat.

† The dynamic range of pairwise distances is the ratio between the largest and the smallest pairwise distances. When this ratio is large, CPM cannot preserve the *large* distances because they convert to small probabilities, that in turn imply small weights in the corresponding terms in the objective function of the optimization, making these terms insignificant.

## Notes

https://github.com/zhiqianZ/supCPM.git

### Competing Interest Statement

The authors have declared no competing interest.

