## Supplementary material for "Supervised Capacity Preserving Mapping: A Clustering Guided Visualization Method for scRNAseq data"

To illustrate the effect of the parameter $d$ in the supCPM algorithm, we simulated a data with three sphere clusters on a straight line with 20 dimensions. From Fig S1, we could find that compared to the Euclidean distance, cluster with geodesic distance is less likely to suffer from the shape deformation. Additionally, the suitable choice of degree of freedom could prevent the stretching of cluster shape. But a overlarge degree of freedom would twist the location of cluster.

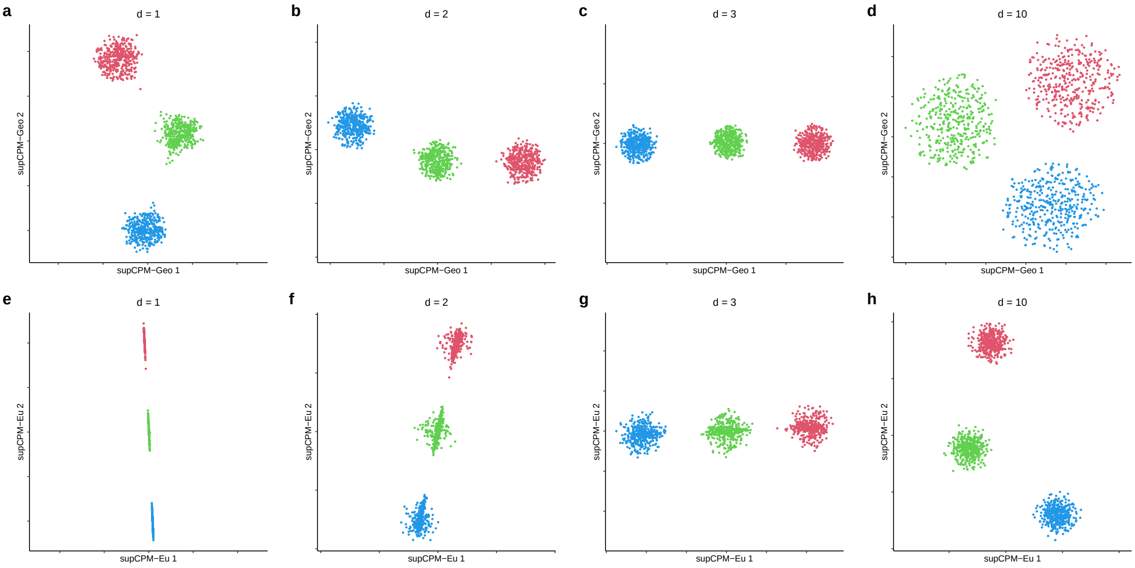


Figure S1: **The influence of the parameter, degree of freedom, $d$.** (**a**)-(**d**) are supCPM with geodesic distance. (**e**)-(**h**) are supCPM with Euclidean distance. From left to right, the degree of freedom takes the value 1, 2, 3 and 10.
